# Integrative Genomic and Regulatory Network Analysis Reveals Adaptive Mechanisms to Salt–Alkalinity Stress in *Brassica fruticulosa*

**DOI:** 10.64898/2026.02.02.703210

**Authors:** Sílvia Busoms, Rhianah Sandean, Glòria Escolà, Kuswati Kuswati, Marek Šlenker, Gabriela Šrámková, Levi Yant, Antoni Garcia-Molina

## Abstract

Salinity poses a widespread and increasing threat to plant fitness, ultimately constraining agricultural productivity worldwide. An inherent roadblock to understanding the precise physiological impacts of high-salinity soils is the frequent co-occurrence of multiple stressors. In calcareous soils, salinity typically coincides with alkalinity. To address this realistic combinatorial stress scenario, we deconstructed the enhanced performance of the coastally distributed, salt-tolerant *Brassica fruticulosa* under salt-alkaline conditions using comparative physiological, transcriptomic, and genomic analyses across major brassica crops. First, to gain a high-resolution genomic view, we generated phased, chromosome-level genome assemblies of *B. fruticulosa* and performed cross-species comparisons of transcriptome-derived Gene Regulatory Networks (GRNs) among important related crop models with contrasting salt tolerances. These results revealed that *B. fruticulosa* mounts predominantly root-centered transcriptional responses to cope with high salinity, whereas salt-sensitive species rely largely on shoot-level mechanisms to mitigate salt toxicity. Consistently, regulatory modules within GRNs diverged substantially between organs and among species, reflecting distinct adaptive programmes of varying efficacy. Functional categorisation of transcription factors with high centrality in *B. fruticulosa* shoot GRNs highlighted processes related to iron (Fe) homeostasis, suggesting that effective maintenance of Fe allocation to aerial tissues supports biomass retention under combined salt and alkalinity stress. Collectively, these findings establish *B. fruticulosa* as a valuable new model for dissecting adaptation to salinity in natural environments and provide mechanistic insight into the regulatory architecture underlying salt–alkaline tolerance.

## Introduction

Soil salinity is a major stressor limiting crop productivity worldwide, affecting more than 20% of irrigated land, and is progressively expanding due to climate change and unsustainable agricultural practices (FAO, 2025). High concentrations of soluble salts in soil impose a two-phase stress on plants: an initial osmotic stress that restricts water uptake, followed by ionic toxicity caused mainly by Na⁺ and Cl⁻ accumulation, which disrupts nutrient balance and cellular homeostasis (Munns, 2002). These primary effects are accompanied by oxidative stress due to excessive reactive oxygen species (ROS) and metabolic imbalances. To manage salinity, plants respond through mechanisms such as ion compartmentalization, osmolyte synthesis (e.g., proline, glycine betaine), activation of antioxidant systems, and hormonal signalling (ABA, ethylene) (Hossain & Dietz, 2016). Nevertheless, prolonged or severe salinity often impairs photosynthesis, growth, and yield, making salt tolerance a key target for crop improvement. Therefore, most studies have focused on simple neutral salt stress, whereas less attention has been paid to salt–alkaline stress, a more realistic condition in calcareous soils such as those widespread across the Mediterranean coasts. This complex stress imposes additional physiological and metabolic constraints, yet its underlying molecular mechanisms remain poorly characterized (Busoms et al. 2025).

Plant adaptation often relies on physiological and metabolic plasticity, allowing dynamic transitions from basal to acclimated states in response to environmental stresses without compromising fitness. These transitions involve reconfigurations of the transcriptome, mediated by gene regulatory networks (GRNs) (Escolà et al., 2026; Jain et al., 2024). Within the Brassicaceae, millennia of domestication have forged diverse vegetables and oilseed species. Despite this long history, domesticates remain vulnerable to abiotic stresses like salinity, drought, and heat, partly due to genetic bottlenecks associated with the domestication process (Cheng et al., 2016; Raza et al., 2020). In contrast, wild relatives offer a valuable source of adaptive diversity, including novel stress tolerance mechanisms that can be introgressed into domesticates (Quesada-Martínez et al., 2021). However, complex traits governed by multiple genes and regulatory networks pose challenges for stable introgression (Katche et al., 2019). These limitations underscore the need for integrative systems-wide approaches to unravel the dynamic interactions underlying stress adaptations.

Recent advances have deepened our understanding of salinity and alkalinity tolerance in Brassicaeae species, highlighting the roles of ion homeostasis, osmolyte accumulation, ROS detoxification, and hormonal signalling (Javid et al., 2012; Almira-Casellas et al., 2024). However, these insights are largely derived from long-established models. Expanding systems-level research to new wild relatives could uncover novel alternative adaptive strategies. Although high-quality genomes are available for cultivated Brassicas (e.g., Chen et al., 2021; Zhang et al., 2025), genomic resources for wild taxa remain incomplete (*GoaT*, 2025), hindering the genomic study of stress tolerance.

To bridge this gap, here we dissect salt–alkaline tolerance in a powerful new model, the wild, salt-alkaline adapted *Brassica fruticulosa* (Busoms et al., 2024), in comparison with major Brassica crops, through an integrative framework combining high-quality genome assembly, comparative transcriptomics, gene regulatory network (GRN) reconstruction, and physiological analyses. We first generate chromosome-level genome assemblies of *B. fruticulosa* enabling comparative analyses within the family. Building on this resource, we investigated how regulatory networks coordinate adaptive responses to combined salinity and alkalinity stress across four Brassicaceae species with contrasting tolerance levels: *B. fruticulosa* and *Brassica napus* (tolerant) versus *Brassica nigra* and *Sinapis alba* (sensitive). Cross-species comparison of GRN topology revealed differential organ specialisation in mounting salinity-responses between tolerant and sensitive species. Functional dissection of GRNs further identified key regulators of iron (Fe) homeostasis as central components sustaining the adaptive response of *B. fruticulosa* to salt–alkali stress. Collectively, our work reveals a novel mechanism underlying agronomically important adaptive variation in a powerful new model system, underscoring the relevance of integrative genomic and network-based approaches to unravel complex adaptive strategies and establishing a foundation for exploiting wild genetic resources in crop improvement.

## Results

### *Brassica fruticulosa* exhibits enhanced tolerance and physiological performance to salinity under alkaline conditions

Tractable models of adaptation to salinity under alkaline conditions remain elusive. Our previous work characterised *B. fruticulosa* as a salt-tolerant species distributed in siliceous and calcareous soils (Busoms et al., 2024). Therefore, we decided to monitor in detail the performance of *B. fruticulosa* under alkalinity and contrast it to other closely-related cruciferous species, namely *B. napus*, a widely cultivated crop with moderate salt tolerance; and *Sinapis alba* and *B. nigra*, as two crop species with clear glycophytic behaviour (Ashraf & McNeilly, 2004). We set up a platform to systematically cultivate plants semi-hydroponically under standard ½ Hoagland Solution (HS) (Control), alkalinity (Alk, 15 mM NaHCO_3,_ pH 8.3), neutral salinity (Salt, 150 mM NaCl), and salinity in combination with alkalinity (Sal-Alk, 15 mM NaHCO_3_ + 135 mM NaCl, pH 8.3) (see methods). Qualitatively, all species under salt-alkalinity exhibited reduced growth, with evident reduced rates of leaf generation. However, this reduction, as well as the decline in turgor state, was most pronounced in *S. alba* and *B. nigra* (**Figure S1**). Consistently, dry biomass and relative water content (RWC) were greatly reduced in these two species under salt-alkalinity compared to controls (**Figure 1A, B**). In contrast, leaf proline levels increased in all species across all treatments, but *B. fruticulosa* and *B. napus* showed a more balanced accumulation pattern, maintaining high levels under combined salt-alkaline stress (**Figure 1C**). Moreover, *B. fruticulosa* and *B. napus* maintained a lower leaf Na:K ratio under both neutral and alkaline salinity compared to the sensitive species (**Figure 1D**), suggesting more efficient ionic homeostasis through selective uptake of K⁺ and reduced accumulation of Na⁺ under saline conditions. These results indicate that under salt-alkalinity *B. fruticulosa* and *B. napus* engage effective stress mitigation strategies, whereas *S. alba* and *B. nigra* exhibit a predominantly damage response.

**Figure 1.**
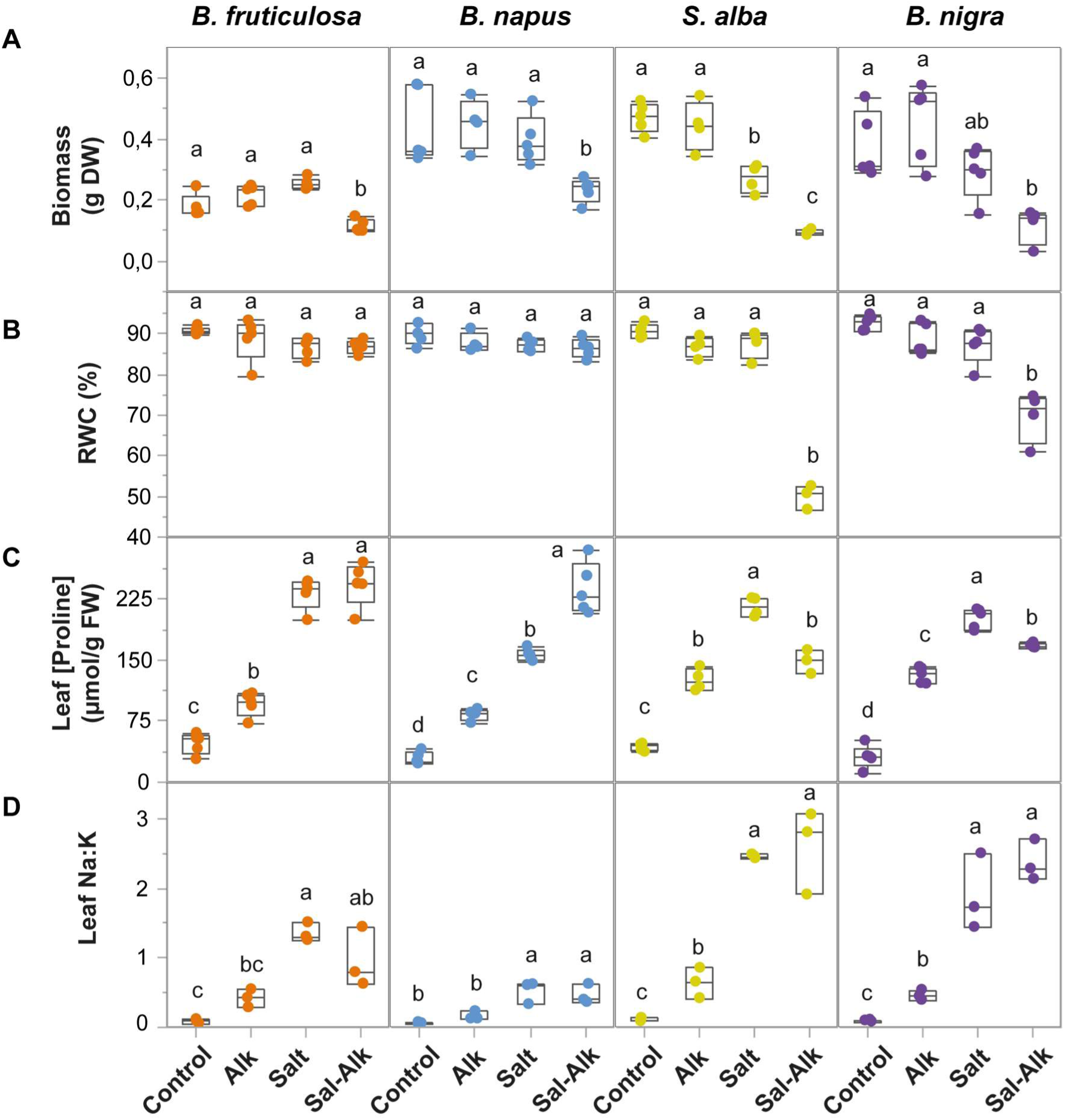
Performance of four Brassicaceae species under salt and alkaline stress. **A**) Biomass (g DW), **B**) relative water content (RWC, %), **C**) leaf proline concentration (umol/g FW), and **D**) leaf Na:K of 25d old *B. fruticulosa*, *B. napus*, *S. alba,* and *B. nigra* plants cultivated under Control (½HS, pH 5.9), Alk (½HS + 15 mM NaHCO_3_, pH 8.3), salinity (Salt; ½HS + 150 mM NaCl, pH 5.9), and salt-alkalinity (Sal-Alk; ½HS + 15 mM NaHCO_3_ + 135 mM NaCl, pH 8.3) conditions for 10 days. Letters indicate significant differences between treatments for each species (adjusted *P* ≤ 0.05, Tukey’s HSD test).

### Chromosome-level genome assembly and annotation of *Brassica fruticulosa*

The relative capacity of *B. fruticulosa* to overcome salt alkalinity led us to pursue a genomic basis for this adaptation. Using an individual from the salt-alkaline-tolerant population ESC, we first estimated genome size by flow cytometry (following Doležel et al., 2007) to be 450 Mb (1C). We then isolated HMW DNA and generated 53 Gb HiFi (∼118× coverage) and 32 Gb HiC (∼71× coverage) data. Using the HiFi data, we confirmed the flow cytometry genome size estimate by K-mer-based analysis to be 454 Mb with 0.6% heterozygosity (K=31; Figure S2A). Primary assemblies with *Hifiasm*, scaffolding with *YaHS*, Hi-C data curation in *Pretext*, and contamination removal with *Blobtools* (see Methods) yielded two chromosome-level, telomere-to-telomere haplomes (two independently assembled representations of the parental chromosome sets), with total lengths of 406 and 407 Mb, respectively (**Figure S2B,C**). Contig N50 was 12 and 16 Mb for each of the two haplomes, and scaffold N50 was 50 and 51 Mb (**Table 1**). Both final chromosome-scale assemblies comprised 8 scaffolds per haplome, the expected number for *B. fruticulosa*.

**Table 1.** Assembly metrics for the *B. fruticulosa* genomes.

| Metric | Haplome 1 | Haplome 2 |
| --- | --- | --- |
| Number of scaffolds | 8 | 8 |
| Total length (bp) | 406,835,166 | 406,312,041 |
| Scaffold N50 (bp) | 49,504,733 | 50,878,404 |
| Largest scaffold (bp) | 59,767,894 | 59,711,205 |
| Contig N50 (bp) | 11,893,649 | 16,350,409 |
| Largest contig (bp) | 32,575,000 | 31,411,000 |
| GC content (%) | 36.04 | 36.03 |

Assembly completeness was very high, with the *Merqury* completeness estimate (Rhie et al., 2020) at 99.3% across both haplomes, with a consensus base quality of QV 68 (∼1 error per ∼7 × 10⁶ bases). Gene space completeness, assessed by analysis of Benchmarking Universal Single-Copy Orthologs (BUSCOs) in *Compleasm* (Huang & Li, 2023), estimated the genome completeness at 99.4% complete for haplome 1 and 99.5% complete for haplome 2. Centromeric satellite repeat analysis indicates metacentric or submetacentric chromosomes, which are similarly positioned in each haplome (**Figure S2D,E**). The haplomes themselves are highly homologous, with no evidence of large-scale structural variation (**Figure S2F**).

Multiple-evidence-guided gene annotation identified 40,450 and 39,545 protein-coding genes in Haplomes 1 and 2, respectively (68,212 and 66,283 gene models including isoforms). The repeat identification pipeline detected 307,621,894 bp and 307,176,606 bp of interspersed repeats and low-complexity DNA sequences in Haplome 1 and Haplome 2 assemblies, representing approximately 75.6% of each haplome. BUSCO analysis indicated high completeness, with 94.3% and 93.0% of complete core genes for Haplomes 1 and 2, respectively. Consistently, OMArk evaluation reported overall completeness of 93.98% for Haplome 1 and 93.33% for Haplome 2, with no detected contaminants and only a small proportion of uncharacterized proteins (approximately 3%). Collectively, these metrics indicate that both *B. fruticulosa* genome assemblies are highly complete, consistently scaffolded, and of good quality, with a reliable gene annotation for comparative genomic and transcriptome analyses.

### Transcriptome responses to salt-alkaline conditions correspond to sensitivity

The completion of high-quality annotations of the *B. fruticulosa* genome next allowed us to infer specific adaptive strategies by deconstructing transcriptional responses in salt-alkali-tolerant *B. fruticulosa* and to perform a cross-species comparison with other crucifers. To do this, we first profiled the transcriptomes from roots and shoots of the four Brassicaceae species across the four treatments. RNA-Seq datasets were filtered for transcripts exhibiting significant differences attributable to alkalinity or salinity treatments (adjusted *P* ≤ 0.05, Likelihood ratio test), or due to the interaction of the two factors (absolute log_2_-fold change ≥ 1, adjusted *P* ≤ 0.05, Wald’s test) (**Dataset S1-S4**).

Overall, salinity imposed the greatest breadth (in terms of numbers of transcripts affected) of transcriptional changes in both roots and shoots, although with different relative degrees depending on the species (**Figure 2A**, **Table 2**). In the more tolerant species, *B. fruticulosa* and *B. napus* exhibited over twice as many significantly altered transcripts in roots than in shoots (**Table 2**). Conversely, glycophytic species *S. alba* and *B. nigra* showed more changes in shoots than in roots (8,402 vs 5,111 and 16,899 vs 10,642, respectively) (**Table 2**). In addition, alkalinity tended to have a similar impact on both roots and shoots in those two species (**Table 2**). These data provide a primary hint towards the importance of roots for counteracting salt stress in tolerant species, whereas salt toxicity ultimately impacts the shoots in the sensitive species. Contrastingly, transcriptome changes due to the interaction between alkalinity and salt were more marked in the glycophytic species, namely in *B. nigra* roots (8,636 transcripts) and in shoots of both *S. alba* and *B. nigra* (4,057-8932 vs 97-120 transcripts) (**Table 2**). These data show that the magnitude of transcriptomic changes under saline conditions correlates with species sensitivity, showing a link between stress responses and gene expression dynamics.

**Figure 2.**
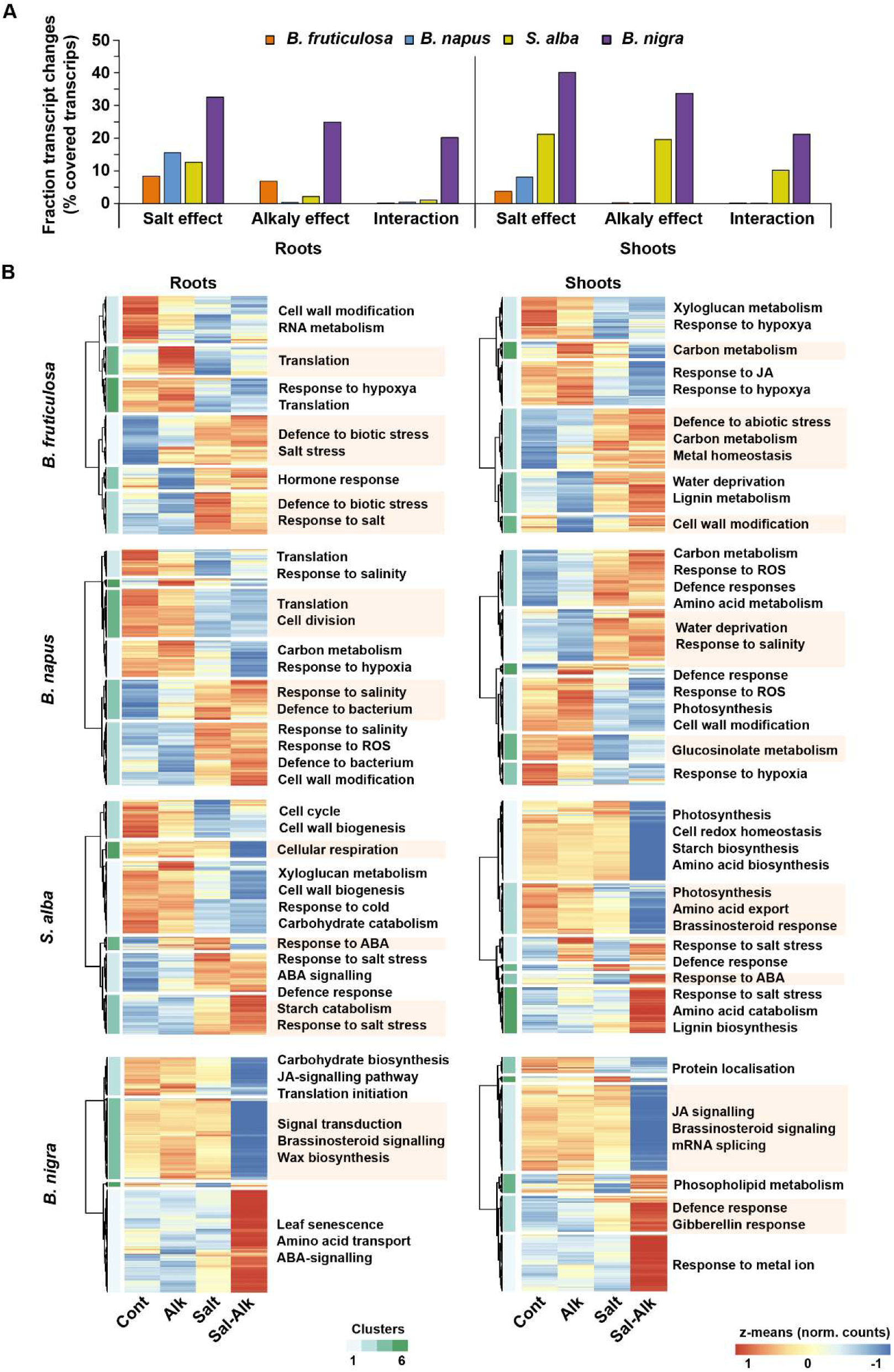
Global transcriptome responses to salt-alkaline conditions correspond to species sensitivity. **A**) RNA-seq data generated from 25-day-old plants cultivated under Control (½HS, pH 5.9), Alk (15 mM NaHCO3, pH 8.3), Salt (150 mM NaCl, pH 5.9), and Sal-Alk (15 mM NaHCO3 + 135 mM NaCl, pH 8.3) conditions. Data were filtered for significant changes in transcripts by salt or alkaline treatments (adjusted P ≤ 0.05, Likelihood ratio test), or the interaction between factors (absolute log2 fold-change ≥ 1, adjusted P ≤ 0.05, Wald’s test). Bar plots represent the percentage of transcriptional changes detected per species in comparison to the global coverage. **B**) Heatmaps were drawn with z-means of normalised counts from RNA-Seq for all significant transcriptome changes per species using the Ward D2 method for hierarchical clustering. Arabidopsis orthologs for genes within clusters were functionally annotated based on significant enrichment in Gene Ontology (GO) terms for biological processes (absolute log2 fold enrichment ≥ 1, adjusted P ≤ 0.05; Fisher’s exact test).

**Table 2.** Transcriptome changes in response to salt-alkalinity treatments. Listed are the number of transcriptional changes determined as indicated in **Figure 3A**.

| Species | Roots |  |  | Shoots |  |  |
| --- | --- | --- | --- | --- | --- | --- |
|  | Salt effect | Alkalinity effect | Interaction | Salt effect | Alkalinity effect | Interaction |
| *B. fruticulosa* | 4,044 | 3,285 | 114 | 1,599 | 123 | 97 |
| *B. napus* | 12,056 | 322 | 380 | 5,903 | 185 | 120 |
| *S. alba* | 5,111 | 879 | 454 | 8,402 | 7,774 | 4,057 |
| *B. nigra* | 13,889 | 10,642 | 8,636 | 16,899 | 14,189 | 8,932 |

Next, we extracted groups of transcripts with correlated expression patterns using hierarchical clustering of dynamic expression values (**Dataset S1–S4**). Clusters were functionally characterised by their significant enrichments in gene ontology (GO) terms for biological processes using cognate Arabidopsis orthologues (**Dataset S5, S6**). Roots of *B. fruticulosa* and *B. napus* displayed a shared trend characterised by a general decrease in transcripts related to cell wall modification, cell division, and translation during salt and salt-alkaline treatments (clusters 2, 5, 6 in *B. fruticulosa*; clusters 1, 2, 5 in *B. napus*). Conversely, the same conditions induced an increase in transcripts responsive to salt and to biotic-stress defence (clusters 1, 3 in *B. fruticulosa*; clusters 3, 4 in *B. napus*) (**Figure 2B**).

A contrasting pattern was evident in shoots, where both *B. fruticulosa* and *B. napus* responded similarly to the salt treatment by decreasing expression of transcripts involved in carbon metabolism and cell wall processes (clusters 1, 2 in *B. fruticulosa*; clusters 2, 4 in *B. napus*) while upregulating those related to carbon catabolism, defence, and salt responses (clusters 3 to 6 in *B. fruticulosa*; clusters 1, 3 in *B. napus*) (**Figure 2B**). In both species, the degree of transcriptome changes tended to be slightly stronger under salt-alkalinity. These expression patterns are consistent with an arrest of central biological processes in roots and enhanced carbon mobilisation in shoots to redirect resources towards sustaining the effective response to stress.

Transcriptome changes in roots of *S. alba* and *B. nigra* showed a distinct pattern in which the salt-alkaline treatment caused substantial perturbations in gene expression, particularly in shoots (**Figure 2B**). There, salt-alkalinity provoked repression of transcripts involved in brassinosteroid signalling (cluster 3 and cluster 2 in *S. alba* and *B. nigra*, respectively). Conversely, transcripts associated with defence responses (cluster 2 and cluster 3 in *S. alba* and *B. nigra*, respectively) commonly increased in shoots of both *S. alba* and *B. nigra* (Figure 3B). Shoots of *S. alba* displayed additional enriched features, namely lower abundance of transcripts for photosynthesis (clusters 1 and 3) and higher abundance of salt- and ABA-responsive genes under salt-alkalinity (**Figure 2B**). In roots, both *S. alba* and *B. nigra* showed—similar to the tolerant species—a decrease in transcripts involved in cell wall biogenesis (clusters 1, 3 in *S. alba*) and in carbon metabolism and translation in *B. nigra* (clusters 2, 3), while ABA-related and salt-responsive transcripts were activated (clusters 3, 4 in *S. alba*; cluster 1 in *B. nigra*) (**Figure 2B**). Collectively, the transcriptomic response associated with salinity stress tolerance reveals conserved features, namely the reconfiguration of translational elements, the remodelling of the cell wall, the activation of defence components and carbon-related metabolic pathways, in addition to salt-responsive elements, though these persist in an organ-dependent manner.

**Figure 3.**
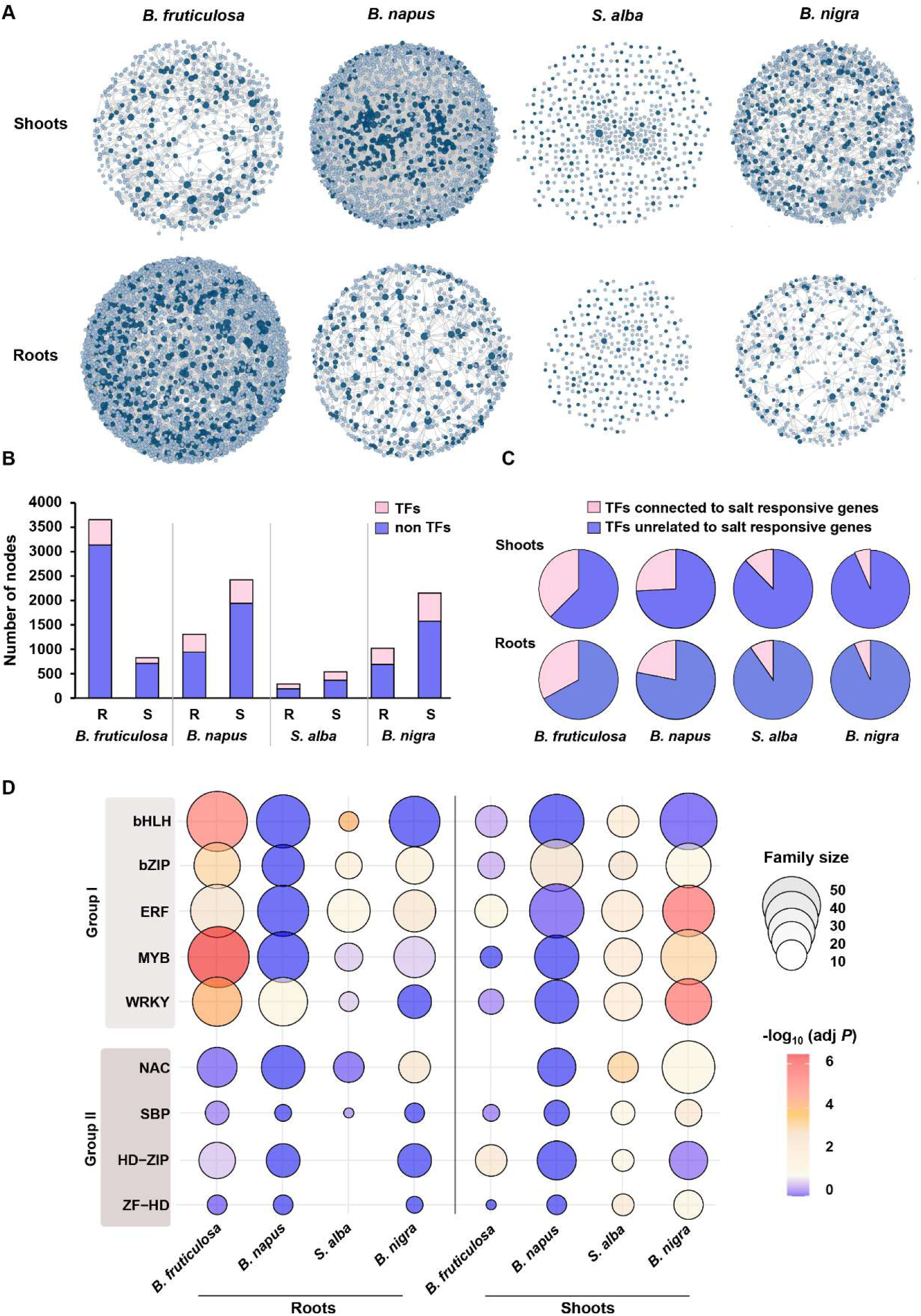
Contrasting organ-specific gene regulatory networks (GRNs) underpin species-specific responses. **A**) GRNs were constructed using RNA-Seq data for TFs and changing transcripts. **B**) Bar plot depicting the scale (number of nodes) of the GRNs and the fraction of TFs included. **C**) Percentage of TFs connected to at least one Arabidopsis orthologue gene related to salt or osmotic stress. **D)** Enrichment of GRNs in families of TFs. The size of the bubble is proportional to the number of TF per family, and the colour code indicates the significance of the enrichment [-log_10_ (adjusted *P*, Fisher’s exact test).

### Opposing prevalence of roots and shoots in the salinity responses of tolerant and sensitive species

To provide a system-wide interpretation of the different responses to salinity among species, we reconstructed Gene Regulatory Networks (GRNs) representing the relations between regulatory modules (transcription factors, TFs) and specific modulated transcripts (targets) (see Methods). GRNs operating in roots and shoots (GRN_roots_, GRN_shoots_) were inferred from RNA-Seq data for transcripts in **Datasets S1–S4** and from the complete set of potential TFs in each crucifer species using the GENIE3 algorithm (see Methods) (**Figure 3A** and **Dataset S7**). A major contrast was immediately evident: topologically, *B. fruticulosa* was the only species for which the scale of the GRN_root_ exceeded that of the GRN_shoot_ (strikingly, by approximately seven-fold), whereas in the remaining species, GRN_shoots_ encompassed twice as many nodes as GRN_roots_ (**Figure 3B**). Thus, we hypothesise that the tolerance acquired by *B. fruticulosa* lies in a dedicated transcriptional reprogramming in the roots.

Across all GRNs, the proportion of nodes corresponding to transcription factors ranged from 15 to 30% (**Figure 3B**). To assess the relevance of TFs in sustaining the response to salt stress, Arabidopsis orthologues of 420 salt- and osmotic-responsive genes were mapped onto each GRN, and the connected TFs were retrieved (**Dataset S7**). Approximately one-third and one-quarter of TFs in the respective GRNs of B*. fruticulosa* and *B. napus* were linked to at least one salt-responsive gene, respectively, whereas in *S. alba* and *B. nigra* GRNs the proportion remained below 10% (**Figure 3B**). This finding suggests that GRNs of tolerant species are more optimally configured to activate specialised components required to withstand salt episodes.

For finer-scale understanding, we next decomposed GRNs in terms of specific TF family composition. The top significantly enriched TF families (adjusted *P* ≤ 0.1, Fisher’s exact test; **Dataset S9**) revealed two main constituents. On one side, TFs belonging to stress-responsive families such as bHLH, bZIP, ERF, MYB, and WRKY displayed antagonistic distribution patterns among species and between GRN_roots_ and GRN_shoots_ (Group I). These TF families were particularly prominent in the *B. fruticulosa* GRN_root_, whereas they were more prominent in the GRN_shoots_ of *S. alba* and *B. nigra* (**Figure 3D**). *B. napus* exhibited enrichment only for the WRKY family (GRN_root_) and bZIP (GRN_shoot_) (**Figure 3D**). This broad TF-family profile indicates that *B. fruticulosa* roots are distinctively prepared to orchestrate the response to salinity, while sensitive species must deploy the counteracting response in their aerial tissues. On the other side, members of the NAC, SBP, HD-ZIP, and ZF-HD families appeared exclusively in the GRN_shoots_ of *S. alba* and *B. nigra* (Group II) (**Figure 3D**). This indicates that salt-sensitive species rely on compensatory TF recruitment in shoots when roots fail to adequately manage environmental salinity.

### Contrasting core salt-responsive regulators between species

Given the asymmetric contribution of roots and shoots to the salinity response, GRNs were further deconstructed to contrast the recruitment of salt- and osmotic stress-responsive genes per organ in each species. Organ-specific composition differed greatly among all four species, with a much larger salt-responsive fraction in *B. fruticulosa* roots than in shoots (133 *vs.* 27; 8 shared; **Figure 4A**), which was relatively balanced in the rest of the species: *B. napus* (69 *vs*. 97; 12 shared), *S. alba* (9 *vs.* 15; none shared), and *B. nigra* (13 *vs.* 26; 4 shared). Overall, there was a broadly species- and organ-specific composition of GRNs in terms of salt-responsive transcripts (**Figure 4B**). Only a minimal overlap existed between *B. fruticulosa* GRN_shoot_ and *B. napus* GRN_shoot_ (15 transcripts) and GRN_root_ sets (13 transcripts). Further comparison to explore potential similarity in TFs targeting Sal-related transcripts across networks also showed that most TFs employed were highly GRN-specific (**Figure 5C**), underscoring the distinctiveness of each species and organ transcriptome.

**Figure 4.**
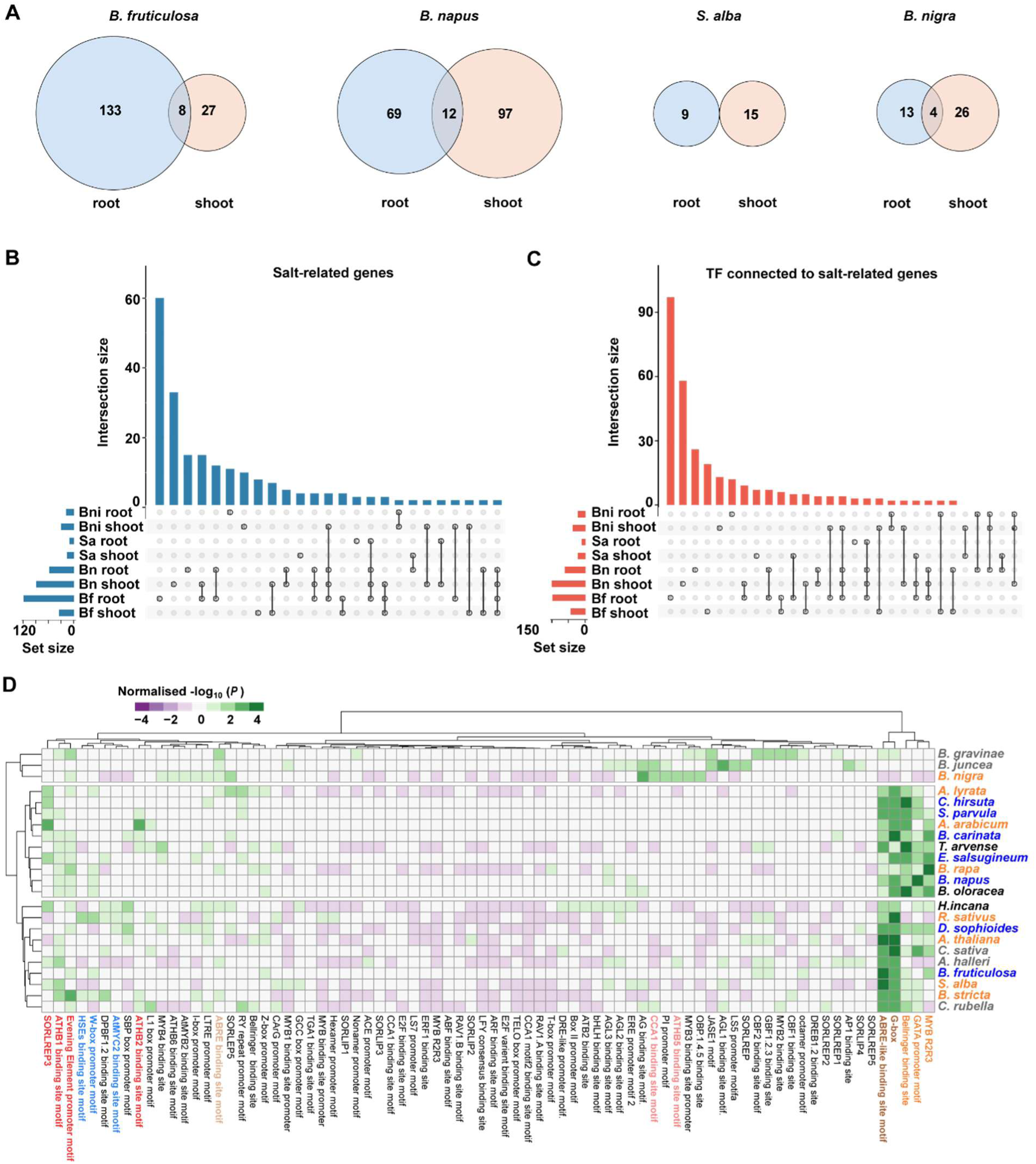
Contrasting salt-responsive regulatory features defining differential adaptive responses. **A**) Overlap in orthologs involved in salt and osmotic stress responses in GRNs of roots and shoots. The size of the fraction is proportional to the number of transcripts included. Multiple comparisons among species and organs (roots, shoots) for the salt-responsive transcripts (**B**) and the TFs connected to them (**C**). **D**) Hierarchical clustering for z-scores of the significance in the presence of putative CREs [-log_10_(*P*), one-tailed Z-test] in the promoters of salt-responsive genes in the indicated species compared to 10,000 randomly selected promoters. Halophyte species are highlighted in blue, glycophytes in orange, moderately tolerant in dark grey, and species not well-characterised in light grey.

**Figure 5.**
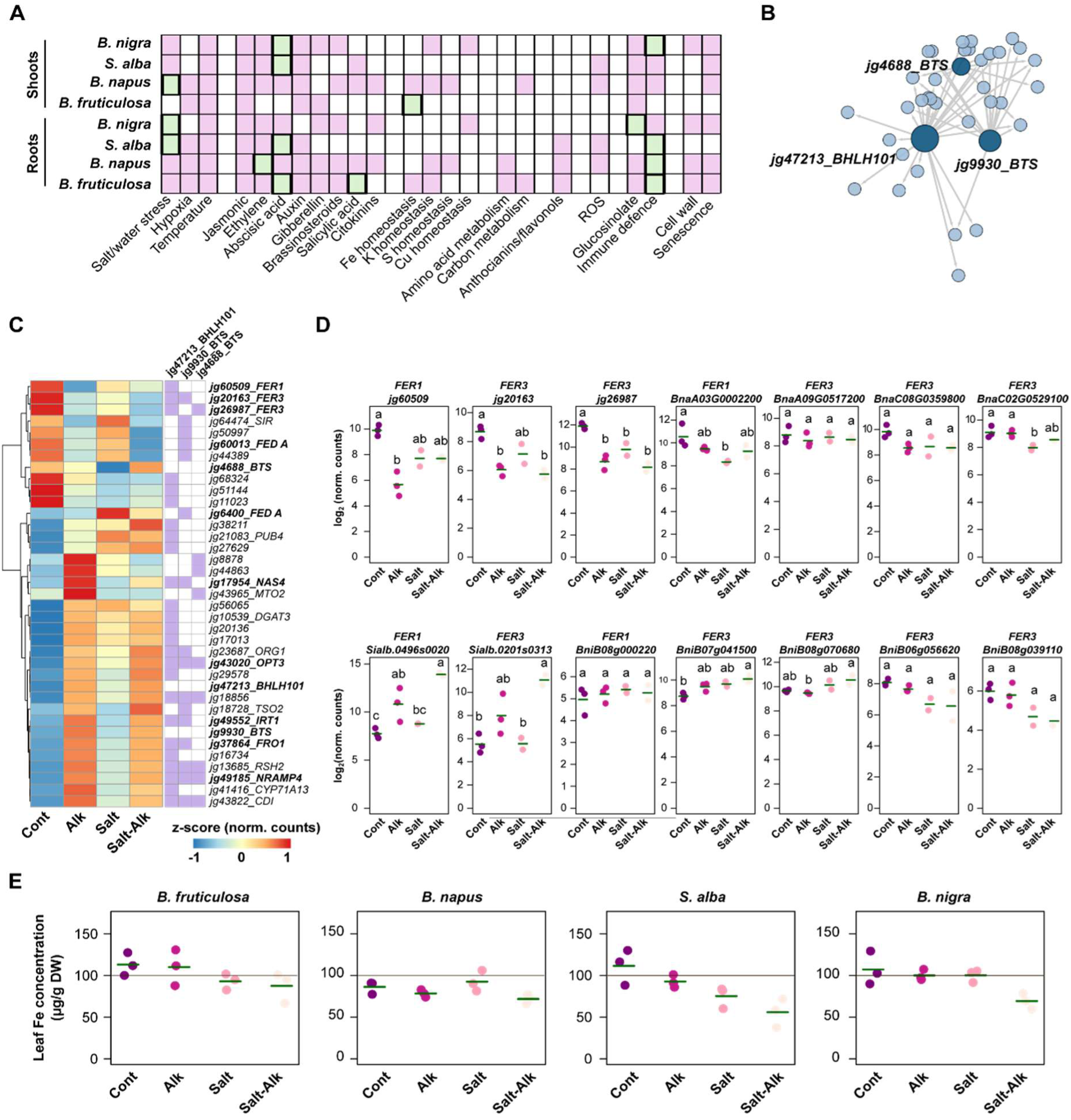
Iron homeostasis as a central component of effective salinity responses. **A**) Summary of GO terms for the TFs included in each GRN. Significantly enriched terms (adjusted P ≤ 0.05, Fisher’s exact test) are indicated in light purple. Terms that include TFs qualifying within the top-five hubs in each GRN are in light green. **B**) Sub-network of Fe-responsive TFs in *B. fruticulosa* shoots sustained by the main hub bHLH101, as well as two BRUTUS (BTS) homologues. **C**) Heatmap with hierarchical clustering according to the Ward D2 algorithm, displaying the changes in z-means of normalised counts for transcript steady-state levels of the features in the subnetwork in **B**. Transcripts related to Fe homeostasis are in bold. **D**) Expression of FER orthologue genes in shoots of the four Brassicaceae species (n = 3 independent plants). Green lines indicate the average normalised counts. **E**) Fe content in shoots of the four species under each treatment (n = 3 independent plants). Statistically significant groups according to Tukey’s post-hoc test (adjusted *P* ≤ 0.05) are indicated with letters.

The contrasting TF deployment across each GRN motivated us to investigate evolutionary dynamics in the regulatory architectures of key salt-related target effector genes. To do this, regulatory regions —2,500 bp upstream to 500 bp downstream— of orthologues of Arabidopsis salt-related transcripts were analysed for differential distribution of 91 consensus TF-binding sites as putative cis-regulatory elements (pCREs) across 23 crucifers —including the four species in this study— with varying tolerances to salinity (**Datasets S8, S10, S11**). For example, *Eutrema salsugineum* and *Schrenkiella parvula* were representative of halophytes, whereas *S. alba*, *Raphanus sativus*, and *Arabidopsis* were representative of glycophytes (**Dataset S11**). The distribution of pCRE enrichments [normalised −log_10_ (*P*), one-tailed Z-test] was used to perform hierarchical clustering, which separated most salt-tolerant (blue labels) and moderately salt-tolerant crucifers (black labels) from glycophytes (orange labels) and three additional species (*B. gravinae*, *B. juncea*, *B. nigra*) (**Figure 4D**).

The glycophyte group was mainly characterised by a marked enrichment of the ABA-responsive element (ABRE)-like motif and the G-box recognised by bHLH TFs—potential targets of salt- and drought-responsive regulators (pCREs in brown, **Figure 4D**). In addition, promoters in this group tended to include other general stress-responsive pCREs such as HEAT SHOCK FACTOR ELEMENTS (HSEs), W-boxes for WRKY TFs, and MYC2- binding sites (pCREs in light blue, **Figure 4D**). The salt-tolerant group also contained the ABRE-like motif and the G-box, but in combination with a BELLRINGER-binding site, a GATA promoter motif, and the putative salt-responsive MYB R2R3 element (pCREs in light orange, **Figure 4D**), as well as several light-regulated elements such as SORLEPs, ATHB-binding sites, and the Evening Element (pCREs in red, **Figure 4D**). Brassicaceae species in the third cluster were not enriched in ABRE-like motifs, but instead showed enrichment for the canonical ABRE motif, together with light-regulated elements including ATHBs and CIRCADIAN CLOCK ASSOCIATED 1 (CCA1) (pCREs in light brown and light red, respectively, **Figure 4D**).

Intriguingly, several species were mis-clustered relative to their known salt-responsive phenotypes: *A. lyrata*, *A. arabicum*, and *B. rapa* clustered with halophytes, whereas *D. sophioides* and *B. fruticulosa* clustered with glycophytes (**Figure 4D**). This observation led us to speculate that the core salinity regulator response in *B. fruticulosa* might be sufficiently robust to obviate the need for the evolution of novel pCREs in the promoters of salt-responsive genes.

### Iron homeostasis signatures emerge as central features in shoots of salt-tolerant species

We next inspected GRNs to identify the prominent pathways underpinning salinity responses based on the functionality of their regulatory modules. Functional annotation of TFs showing significant GO term enrichment (adjusted *P* ≤ 0.05, Fisher’s Exact test) revealed GRNs involved in: (i) salt and/or water stress, hypoxia, and temperature responses; (ii) hormone responses; (iii) mineral nutrition; (iv) primary metabolism (amino acids and carbohydrates); (v) secondary metabolism (anthocyanins and flavonols); (vi) responses to reactive oxygen species (ROS); (vii) glucosinolates and defence against biotic stressors; and (viii) cell wall remodelling and senescence, depending on the network (**Figure 5A**). The TFs in GRN_root_ of all four species were linked to salt and/or water stress and temperature responses, as well as signalling by jasmonic acid (JA), ethylene, and ABA—major mediators of salt-stress responses—and to defence components (**Figure 5A**). This again underscores the centrality of roots in overcoming salinity. All GRN_shoot_ included TFs associated with temperature responses, JA and auxin signalling, and immunity (**Figure 5A**). Interestingly, the modules of all GRN_shoot_ except that of *B. fruticulosa* partially mirrored the pattern of GRN_root_, with features related to salt responses, ethylene and ABA signalling, and senescence (**Figure 5A**). Consequently, *B. fruticulosa* appears able to articulate an alternative transcriptional programme in shoots during periods of salt stress, in contrast to the other species—which, particularly the glycophytes—rely on a strictly salt-responsive configuration similar to roots.

To pinpoint the key regulatory modules driving the response to salt in each species, we searched for TFs included in enriched functional terms that also qualified as top-five hubs within each GRN (**Dataset S12, Figure 5A**, light green). TF hubs in GRN_root_ overlapped only in defensive features (glucosinolates or immune defence). In addition, TF hubs in GRN_root_ of the tolerant species were related to hormones (ABA and salicylic acid in *B. fruticulosa*; ethylene in *B. napus*), whereas hubs in the two glycophytes were located in salt and/or water stress and defence, consistent with their higher sensitivity (**Figure 5A**). Similarly, TF hubs in GRN_shoot_ of *S. alba* and *B. nigra* were associated with ABA (**Figure 5A**). The main TF hub in GRN_shoot_ of *B. napus* was linked to salt and/or water stress, suggesting that—even in a tolerant species—activation of salt -responsive features in shoots remains necessary to withstand the stress (**Figure 5A**). In GRN_shoot_ of *B. fruticulosa*, an ortholog of the Arabidopsis *bHLH101*—a central regulator of the Fe starvation response—was positioned as a major hub (**Figure 5A**). A closer inspection of the *B. fruticulosa* GRN_shoot_ revealed the presence of two orthologs of the Fe regulator *BRUTUS* (*BTS*), which converged on several bHLH101 targets (**Figure 5B**). Notably, these TFs were connected to Fe-responsive genes such as *FERRITIN1/3* (*FER1/3*), *OPT3*, *IRT1*, *FRO1*, *NAS4*, *NRAMP4*, and *FEDA*. Taken together, our data uncover a potential contribution of Fe-homeostasis features to sustaining salinity tolerance in *B. fruticulosa* shoots.

To evaluate the role of Fe under salt stress conditions in *B. fruticulosa*, shoot transcriptomes were examined specifically with respect to Fe-responsive target genes. As shown in **Figure 5C**, alkalinity —a treatment that reduces Fe bioavailability— triggered a dramatic downregulation of three Fe-storage proteins (*FERs*) compared with controls, a trend that was also maintained under salt and salt-alkalinity. A similar pattern was observed for the transcript of the Fe-containing protein FEDA (**Figure 5C**). Conversely, genes involved in Fe uptake (*FRO1, IRT1, NRAMP4*) and distribution (*OPT3*) remained upregulated under alkalinity, and also under salt and salt-alkalinity (**Figure 5C**). Transcript levels of the Fe-chelating intermediate NAS4 increased under alkalinity but not under salt (**Figure 5C**). Together, these findings indicate that salinity treatments impose the mobilisation and circulation of free Fe pools in *B. fruticulosa* shoots.

To determine whether signalling for Fe mobilisation also occurs in the other species, the expression values of FER orthologs were examined. *B. napus FER1* and one *FER3* (BnC08G0359800) followed dynamics like those observed in *B. fruticulosa* (**Figure 5D**). Surprisingly, *FER* transcripts in *S. alba* tended to accumulate under salt-alkalinity, while those in *B. nigra* showed mixed tendencies, with two *FER3* genes decreasing under salt and the other two remained unaffected (**Figure 5D**). Thus, the tolerant species appear to reallocate Fe to the shoot in response to salinity. Indeed, Fe concentration *in B. fruticulosa* and *B. napus* shoots remained constant across the four treatments (approx. 100–110 and 90 µg/g DW, respectively), within the sufficiency range (100–120 µg/g DW) (**Figure 5E**). In contrast, both salt-sensitive drop to deficiency levels in shoot Fe in salinity (from approx. 110 µg/g DW in the control to 50–60 µg/g DW under salt-alkalinity) (**Figure 5E**). In conclusion, the tolerant species can maintain Fe allocation to shoots as part of their strategy to counteract the imbalances caused by salinity.

### *Brassica fruticulosa* fitness under salt-alkaline stress partly depends on iron sufficiency

Based on the ability of tolerant species to maintain Fe sufficiency in shoots under salt-alkali, we evaluated the performance of two species with strongly contrasting phenotypes (*B. fruticulosa* and *S. alba*) in relation to Fe availability. We contrasted plants grown under control conditions to those cultivated in salt-alkali conditions, combined also with either standard or low Fe supply (50 µM vs. 10 µM; Fe_50_, Fe_10_). Under control conditions, both species displayed normal growth (**Figure 6A**), with comparable biomass (FW), water content (WC: FW/DW), and effective quantum yield of photosystem II (Φ_II_), regardless of Fe availability (**Figure 6B–D**). This indicates that *B. fruticulosa* and *S. alba* can efficiently tolerate moderate Fe limitation.

**Figure 6.**
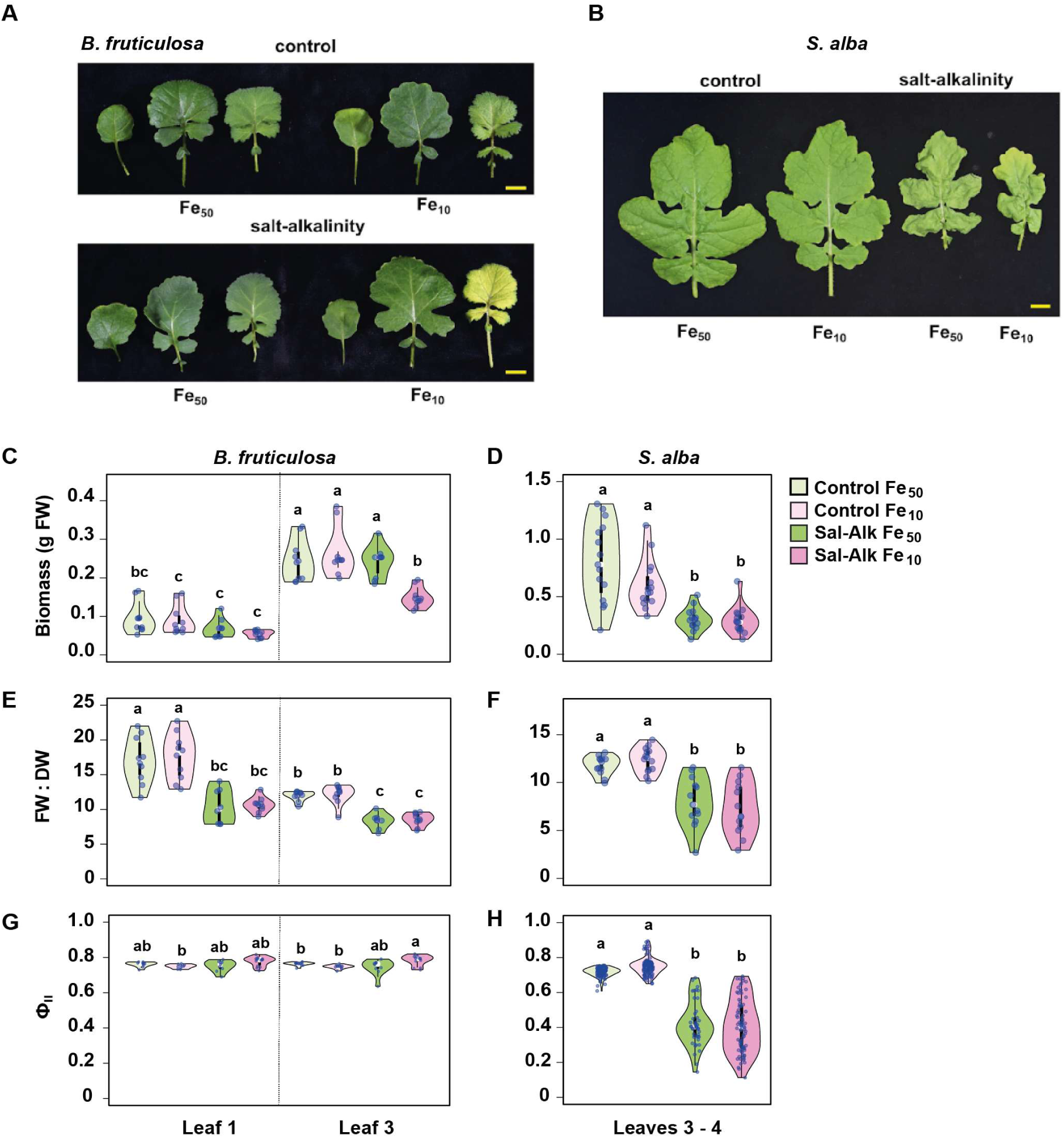
Partial iron dependence of *B. fruticulosa* for adaptation to saline–alkaline conditions. Representative images of *B. fruticulosa* leaves 1, 3, and 5 (**A**) and *S. alba* leaves 3 and 4 (**B**) from plants grown under control or salt-alkaline conditions with sufficient or low Fe (Fe_50_, Fe_10_). Biomass (FW; **C**, **D**), water content (FW/DW; **E**, **F**), and the effective quantum yield of photosystem II (Φ_II_; **G**, **H**) were quantified in the corresponding leaves shown in panels A–B. Violin plots represent data from n = 3 independent experiments. Statistically significant groups according to Tukey’s post-hoc test (adjusted *P* ≤ 0.05) are indicated by different letters.

In *B. fruticulosa*, salt-alkalinity under Fe_50_ did not limit FW in leaves 1 (oldest) and 3 (intermediate), although WC decreased by approximately twofold compared with controls (**Figure 6B–D**). Under Fe_10_, however, salt-alkalinity caused leaf curling in older leaves and chlorosis in younger ones (**Figure 6A**), along with reduced FW in leaf 1 and, more markedly, in leaf 3 (approx. 40%), while WC decreased to levels similar to those in Fe_50_ (**Figure 6B–C**). Nevertheless, Φ_II_ remained unaltered under salt-alkalinity, independent of Fe supply (**Figure 6D**). In contrast, *S. alba* exhibited stunting under salt-alkalinity, with significant reductions in FW, WC, and Φ_II_ (approx. 30–50%) in intermediate leaves (leaves 3–4) under both Fe conditions compared with the respective controls (**Figure 6A–C**). Overall, these results show that although *B. fruticulosa* can withstand salt-alkaline stress even under low Fe, maintaining leaf water status and an intact photosynthetic apparatus, adequate Fe availability remains essential to sustain proper leaf biomass and prevent morphological constraints. Conversely, Fe levels do not modulate the sensitivity to salinity in *S. alba*.

### Conserved hub transcriptional regulators in Brassicaceae species

Since the partial Fe dependence of *B. fruticulosa* in overcoming salt–alkaline stress validated the ability of our model to delineate novel features involved in stress tolerance, we next explored the translational potential of the identified candidate genes in other species. To this end, the TF hubs linked to the most representative biological functions of the regulatory modules in the GRNs of the two salt–alkaline tolerant species *B. fruticulosa* and *B. napus* (**Figure 5A, Dataset S12**) were inspected. Synteny relationships across *B. fruticulosa*, *S. alba*, *B. nigra*, and *B. napus* reveal that these TF hubs are embedded within conserved genomic blocks, supporting their evolutionary stability and functional relevance (**Figure 7**). As previously stated, besides the Arabidopsis orthologue for bHLH101 (AT5G04150) in *B. fruticulosa*, the majority of identified TF hubs -MYB122 (AT1G74080), HBI1 (AT2G18300), BZIP45 (AT3G12250), BZIP53 (AT3G62420), ERF040 (AT5G25810), SDIR1 (AT3G55530), and DPBF3/ABI5-like 2 (AT3G56850)- were associated with stress and hormone signalling, suggesting tight integration of hormonal control within the GRNs. The presence of NAI1 (AT2G22770), involved in ER body development, and a C2H2 BIRD-type transcription factor (AT2G02080) further points to coordinated regulation of cellular organisation and stress-responsive gene expression. Together, the conservation and high connectivity of these hub genes in tolerant species point to a core regulatory module that represents a promising target for transferring salt–alkaline tolerance to more sensitive Brassicaceae species.

**Figure 7.**
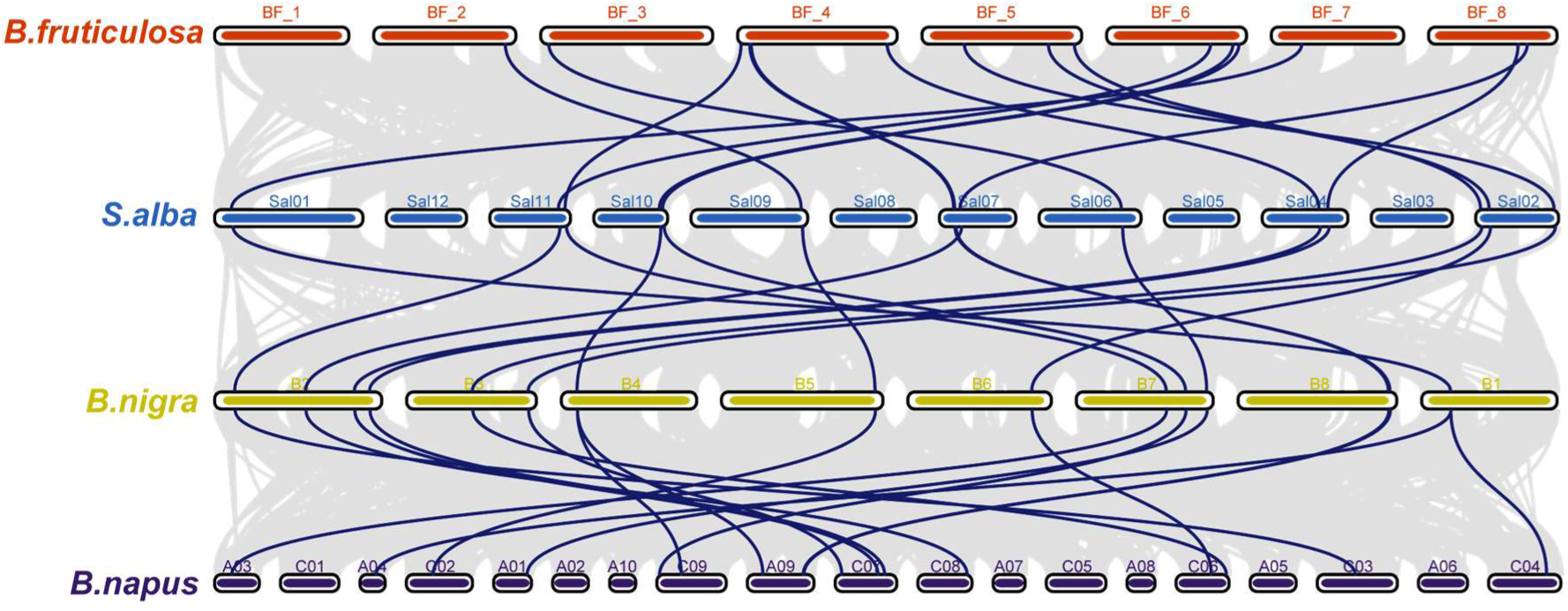
Comparative genomic analysis of syntenic blocks in *B. fruticulosa, S. alba, B. nigra,* and *B. napus*. Syntenic blocks are indicated in grey, and the selected transcription factors acting as hubs in the Gene Regulatory Networks identified in Figure 5A and **Dataset S12** are indicated in blue.

## Discussion

*Brassica fruticulosa* displays pronounced tolerance and superior physiological performance under salt–alkaline stress relative to the other Brassicaceae species here tested, indicating the presence of robust and likely multi-layered adaptive mechanisms. Interestingly, this resilience does not stem from canonical Na⁺ extrusion pathways, as shoot Na:K ratios increased under both salt and salt–alkali treatments (**Figure 1D**). This observation suggests that *B. fruticulosa* relies on alternative strategies beyond simple ion exclusion, potentially involving metabolic reconfiguration, enhanced detoxification capacity, and/or cellular homeostasis programs. Accordingly, *B. fruticulosa* represents a valuable new model to dissect complex, natural adaptations to salinity prevalent in calcareous habitats, while offering promising traits for breeding tolerant Brassica crops suited for saline and alkaline soils.

The generation of quality genome assemblies and annotations for *B. fruticulosa* enabled a comprehensive reconstruction of salt- and alkali-responsive transcriptomes and GRNs, followed by cross-species comparisons with both tolerant and sensitive crucifers. This adaptive GRN topology highlighted a disproportionately large regulatory contribution from roots, particularly in *B. fruticulosa*, compared with *B. napus*. Both tolerant species showed prominent recruitment of WRKY and ZIP transcription factors, well-established regulators of salinity and osmotic stress responses (Hussain et al., 2021; Sun et al., 2025). However, *B. fruticulosa* additionally mobilised other stress-associated TF families, suggesting the presence of a reinforced and more specialised regulatory program to mitigate salinity stress. In contrast, shoots of tolerant species exhibited limited TF enrichment, consistent with the mild toxicity observed in aerial tissues under the applied treatments.

Sensitive species presented an opposite regulatory configuration: their GRN_shoots_, but not GRN_roots_, were enriched in stress-responsive TFs, mirroring physiological phenotypes in which shoots incur substantial damage. Thus, while tolerant *B. fruticulosa* activates root-centric programs that restrict Na⁺-induced injury, sensitive species appear to mount reactive rather than preventive responses, consistent with a survival- rather than tolerance-oriented strategy (He et al., 2017). Comparative GRN analyses further revealed striking species-specific regulatory modules. Although the biological processes associated with the recruited TFs were broadly conserved across species, the identities and network positions of key regulators differed, underscoring lineage-specific solutions to salinity stress. Whether these differences reflect variation in evolutionary history, ecological niche, or inherent efficiency of adaptation to high-pH environments will require further investigation.

Of particular interest was the unique enrichment of Fe homeostasis regulators in the *B. fruticulosa* GRN_shoot_, a feature absent in the other species. Coupled with physiological measurements, these transcriptomic signatures delineate a second axis of divergence between tolerant and sensitive Brassicaceae. Tolerant species maintained shoot Fe concentrations within sufficiency ranges under both salt and salt–alkaline stress, whereas sensitive species, especially under alkaline conditions, displayed pronounced Fe deficiency (**Figure 6**). This suggests that the ability to secure and mobilise Fe to aerial tissues under stress is integral to tolerance as observed in Arabidopsis (Almira-Casellas et al., 2024) and non-Brassicaceae species (e.g., Wang et al., 2023). Indeed, the application of exogenous EDDHA/Fe in rice and Fe nanoenzymes in wheat enhanced growth, nitrogen metabolism, photosynthesis, and transcriptomic changes, markedly improving their tolerance to saline-alkaline stress (Gao et al., 2024; Chen et al. 2025).

Plants in calcareous soils face intrinsic Fe limitation due to reduced Fe bioavailability at high pH. Brassicaceae have long been classified as Strategy I plants, acquiring Fe via ferric chelate reduction mediated by FROs and uptake through IRT1 (Jeong & Connolly, 2009). However, recent findings show that Arabidopsis secretes Fe-mobilising coumarins and subsequently absorbs Fe complexes through alternative transporters under neutral-to-alkaline conditions, a mode resembling Strategy II components (Robe et al., 2025). These discoveries raise the possibility that *B. fruticulosa* may employ an expanded or modified Fe-acquisition repertoire, consistent with its superior Fe homeostasis under salt–alkalinity. Maintaining adequate Fe is advantageous under salinity, as Fe is essential for antioxidant enzymes (superoxide dismutases, catalases, peroxidases) and for sustaining photosynthetic electron transport. Insufficient Fe compromises chloroplast function, but excess Fe can exacerbate oxidative stress through Fenton chemistry (Rout & Sahoo, 2015). Thus, shoot Fe accumulation in tolerant species may help buffer stress-induced ROS while supporting energy metabolism. Conversely, sensitive species may experience compounded stress due to simultaneous Na⁺ toxicity and Fe deficiency.

Our physiological assays reinforce this model. Under salt–alkaline stress, Fe limitation impaired biomass accumulation and leaf morphology in *B. fruticulosa*, although photochemical efficiency (Φ_II_) remained stable even under low Fe. This indicates that Fe contributes significantly, but not exclusively, to salt–alkaline resilience. *Brassica fruticulosa* can maintain photosynthetic function even under low Fe conditions, albeit at the cost of reduced growth, illustrating a trade-off between metabolic protection and biomass production. By contrast, *S. alba* showed no improvement under higher Fe supply, suggesting that Fe availability is not a limiting factor and that its sensitivity arises from different physiological bottlenecks.

Overall, our findings reveal that *B. fruticulosa* employs an integrated strategy combining reinforced root regulatory programs, minimized shoot stress signalling, and effective Fe acquisition and mobilisation, collectively enabling superior tolerance to salt and salt–alkaline conditions. These mechanistic insights establish a foundation for identifying genetic determinants of salinity resilience and for engineering Brassica crops adapted to the increasingly prevalent stress scene of alkaline and saline soils.

## Material and Methods

### Genome assembly and annotation of *Brassica fruticulosa*

High-molecular-weight DNA was extracted from a young seedling of *Brassica fruticulosa* accession ‘ESC’ (L’Escala; Busoms et al., 2024) following a modified CTAB protocol (Doyle & Doyle, 1987) and used for PacBio HiFi and Hi-C sequencing. HiFi reads were generated on the PacBio Sequel Revio platform, and Hi-C libraries prepared from leaf tissue using the Arima Genomics® protocol were sequenced on an Illumina NovaSeq 6000. Genome size was estimated using k-mer distribution analyses (*KMC*; Kokot et al., 2017; *GenomeScope v2.0*; Ranallo-Benavidez et al., 2020) and validated by flow cytometry (Doležel et al., 2007).

A phased, chromosome-level genome assembly was generated using *hifiasm* (Cheng et al., 2021) by integrating HiFi and Hi-C data, followed by scaffolding with *YaHs* (Zhou et al., 2023). Potential contamination was assessed using *BlobTools* (Laetsch & Blaxter, 2017), incorporating gene completeness metrics from *Compleasm* (Huang & Li, 2023) and taxonomic assignments based on *Diamond BLAST* (Buchfink et al., 2021). Assemblies were manually curated using *PretextView* following established rapid curation pipelines, with telomeric and centromeric features validated using *tidk* (Brown et al., 2025) and *TRASH* (Wlodzimierz et al., 2023). Assembly quality and completeness were evaluated using gfastats (Cabanettes & Klopp, 2018) and BUSCO analyses implemented in *Compleasm* (Huang & Li, 2023).

Repetitive elements were identified using a combination of de novo and homology-based approaches. The combined repeat library was clustered to generate a non-redundant repeat dataset, and then, used to mask the *B. fruticulosa* genome using *RepeatMasker* (Flynn et al., 2020). Protein-coding genes were predicted on the repeat-masked genome using an evidence-guided strategy integrating ab initio, homology-based, and transcriptome-based approaches, utilising RNA-seq data from *B. fruticulosa* samples, protein sequences from several related Brassicaceae species, and the OrthoDB12 Viridiplantae dataset (Tegenfeldt et al., 2025). Gene models generated by all three approaches were integrated using *EvidenceModeler* (Haas et al., 2008) and further refined with *PASA* (Campbell et al., 2006) to add untranslated regions and alternative splicing isoforms. Final gene models were filtered to remove entries with internal stop codons, missing introns, or duplicated features. Annotation completeness was evaluated using *BUSCO* (Manni et al., 2021) and *OMArk* (Nevers et al., 2025).

Full methodological details are provided in **Supplementary Methods**.

### Co-linearity and homology inference

Synteny analysis of candidate genes was performed using the One-step MCScanX plugin in *TBtools-II* v.2.3.76 (Chen et al., 2023). Pairwise syntenic relationships among *B. napus, B. nigra, S. alba,* and *B. fruticulosa* were identified using an E-value cut-off of E-value ≤ 1 × 10^−5^ and visualised using the dual synteny plotter module in TBtools-II.

Potential functional orthologs in *B.fruticulosa* were identified using a comparative genomics approach anchored to *B.napus*. Homologous genes were surveyed across *B.nigra, S.alba* and *B.fruticulosa*. Homolog identification was based on sequence similarity and genomic context (location and collinearity). Pairwise genome comparisons were generated using the *JCVI toolkit* v1.5.6 (Tang et al., 2024), which performs whole genome alignments and subsequently filters the results to remove tandem duplications and low confidence matches. A C-score of 0.99 was applied to retain high-confidence matches. The filtered alignments outputs (last.filtered) were used to identify the best homologous gene pairs from the *B.napus* genome across *B.nigra, S.alba,* and *B.fruticulosa*. Collinearity was visualised using the *JCVI toolkit* with a c-score threshold of 0.99.

### Plant materials and cultivation conditions

Seeds of *B. fruticulosa* were collected from Tossa de Mar (TOS population, Busoms et al., 2024) in spring 2022. Commercial seeds of *B. napus* (Grup Morera, Les Preses, Girona, Spain), *S. alba* (Semillas Batlle, Molins de Rei, Barcelona, Spain), and *B. nigra* (Flors Catalunya, Vilassar de Mar, Spain) were used. Seeds were surface sterilised in 10% (v/v) commercial bleach and 0.01% (v/v) Tween-20 for 15 min, followed by several rinses with water. Seeds were stratified for 2 days at 4 °C and sown in a humidified soil:sand:perlite (1:1:1) substrate. Plants were grown in FitoClima 1200 PLH LED cabinets (Aralab, Rio de Mouro, Portugal) under a 10-h light (25 °C; 150 µmol m^−2^ s^−1^)/12-h dark (20 °C) cycle at 40% relative humidity. Plants were irrigated with ½-strength Hoagland solution (pH 5.9) for one week before treatments were applied. The treatments consisted of: control: ½ Hoagland (pH 5.9); alkalinity: gradual irrigation with 5 mM NaHCO₃ (pH 8.3) up to 15 mM over 8 days; salinity: gradual irrigation with 50 mM NaCl (pH 5.9) up to 150 mM over 8 days; and salt-alkalinity: gradual irrigation with 10 mM NaHCO₃ + 40 mM NaCl (pH 8.3) up to 30 mM NaHCO₃ + 120 mM NaCl over 8 days.

To evaluate the effect of Fe on salt–alkaline responses, *B. fruticulosa* and *S. alba* plants were irrigated as described above using standard ½ Hoagland (50 µM Fe-EDTA) or reduced iron (10 µM Fe-EDTA) (Fe_50_ and Fe_10_), reaching maximum concentrations of 20 mM NaHCO₃ + 80 mM NaCl for *S. alba* and 30 mM NaHCO₃ + 120 mM NaCl for *B. fruticulosa*. Plants (25 days old) were harvested and immediately flash-frozen in liquid nitrogen. Experiments were performed in triplicate using pools of at least three plants.

### Determination of physiological parameters and ionomic analysis

Leaf relative water content (RWC) was calculated using fresh weight, turgid weight (after 4 h in water), and dry weight, following the formula: [(FW−DW)/(TW−DW)]×100.

Proline concentration was determined colorimetrically following Bates et al. (1973) with minor modifications. Frozen leaf tissue (50 mg) was homogenised in 3% (w/v) sulfosalicylic acid, and the supernatant was incubated with acid ninhydrin at 98 °C for 60 min. After cooling, toluene was added, and the absorbance of the organic phase was measured at 520 nm. Proline concentration was calculated using a standard curve.

The effective quantum yield of photosystem II (ΦII) was measured on plants dark-adapted for 15 min using an IMAGING-PAM M-Series instrument (Walz, Effeltrich, Germany). Minimal (F₀) and maximal (Fₘ) fluorescence were measured using a saturation pulse (0.5 s; 2700 µmol photons m^−2^ s^−1^; 450 nm), and ΦII was calculated as (F_m_’ – F_o_’)/F_m_’.

For ion composition, ∼0.1 g of dried leaf tissue was digested in 65% (v/v) HNO₃ at 110 °C for 5 h. K, Na, and Fe concentrations were determined by ICP-OES (Thermo Jarrell-Ash, model 61E Polyscan, UK).

### Transcriptome profiling and analyses

Total RNA from roots and shoots (n = 3 independent plants) was extracted using the Maxwell® RSC Plant RNA Kit (Promega, Wisconsin, USA). RNA concentration was measured with a Qubit 2.0 fluorometer (Life Technologies, Carlsbad, CA, USA). Paired-end RNA-Seq libraries (2×150 bp) were prepared and sequenced by Novogene (China) following Illumina protocols.

RNA-Seq analyses were performed on the Galaxy platform (The Galaxy Community, 2024). Reads were trimmed with Trimmomatic and mapped to the corresponding reference genomes using *STAR* (Dobin et al., 2013) (genome versions listed in **Dataset S11**). Read counts were obtained using *featureCounts* (Liao et al., 2014) and analysed with *DESeq2* (Love et al., 2014) according to models testing transcriptional changes due to “salinity”, “alkalinity” (adjusted *P* ≤ 0.05; Likelihood ratio test), or their interaction [absolute log₂(fold-change) ≥ 1, adjusted *P* ≤ 0.05; Wald test].

Gene regulatory networks (GRNs) were inferred from the RNA-Seq datasets using normalised expression values of responsive transcripts and their Arabidopsis orthologues annotated as transcription factors (TFs) in the *Plant Transcription Factor Database* (TFDB 5.0) (https://planttfdb.gao-lab.org/index.php). Networks were inferred using GENIE3 (Huynh-Thu & Geurts, 2018) in RStudio with default parameters. Next, the function *fct_edge_testing()* in the *RStudio* package implemented for the *Dashboard for the Inference and Analysis of Networks from Expression data* (DIANE) (Cassan et al., 2021) was applied to filter edges to obtain networks at different ranges of density (10^−5^ to 10^−2^). A threshold was selected at the density yielding a saturated number of nodes and TFs with more than 10 connections. GRNs were exported and visualised in Cytoscape (Shannon et al., 2003).

### Bioinformatic analyses

Arabidopsis orthologues for the selected brassica species were inferred according to curated databases such as *PLAZA dicots v.5.1* (*Aethionema arabicum, A. lyrata, B. carinata, B. oleracea, B. rapa, C. hirsuta, E. salsugineum*, and *S. parvula*) or the *Brassica napus multi-omics Information Resource* (BnIR) (*B. napus*) (Van Bel et al., 2022); Yang et al., 2023), or computed with the *Orthofinder v.2.5.5* tool (Emms & Kelly, 2019) (*A. halleri, B. fruticulosa, B. gravinae, B. juncea, B. nigra, Camelina sativa, Capsella rubella, Descurinia sophoides, Hirschfeldia incana, Raphanus sativus, S. alba*, and *Thlaspi arvense*) integrated in the Galaxy platform (usegalaxy.eu) with the versions of reference proteomes listed in **Dataset S11**.

Heatmaps were generated from z-scored normalised counts using the *pheatmap()* R package. Hierarchical clustering used the *Ward D2* algorithm, and optimal cluster partitioning was determined by k-means.

GO term enrichment analyses in biological processes [log_2_ (fold-enrichment) ≥ 1, adjusted *P* ≤ 0.05; Fisher’s exact test] were performed with Panther (https://pantherdb.org) using Arabidopsis orthologs. TF enrichments in GRNs were assessed by comparing observed TF frequencies with expectations based on the total number of orthologues per family (adjusted *P* ≤ 0.05). Bubble plots were generated using *ggpubr*.

Multiple comparisons of transcripts and TFs were visualised using Venn diagrams (Venny v2.1) or UpSet plots (R package UpSetR).

For enrichment in promoter cis-regulatory elements (pCREs), promoter regions (-2500 to +500 bp relative to the start codon) were screened for documented Arabidopsis pCREs (**Dataset S10**). The frequency of pCREs was compared to that of 10,000 random genes (P ≤ 0.05; one-tailed Z-test). Heatmaps were generated using z-scores of -log_10_(*P*), and pCRE and species clustering followed the Ward D2 algorithm.

## Supporting information

Dataset S1. Transcriptome changes in Brassica fruticulosa.

Dataset S2. Transcriptome changes in Brassica napus.

Dataset S3. Transcriptome changes in Sinapis alba.

Dataset S4. Transcriptome changes in Brassica nigra.

Dataset S5. Arabidopsis ortholog conversion of transcripts of Brassicaceae species.

Dataset S6. Functional annotation of genes included in the heatmap for transcriptome changes.

Dataset S7. Gene Regulatory Networks.

Dataset S8. Orthologues for salt- and osmotic stress-responsive genes in distinct Brassicaceae species.

Dataset S9. Enrichment of Gene Regulatory Networks in transcription families.

Dataset S10. Potential cis-regulatory elements used in this study.

Dataset S11. List of additional Brassicaceae species used in this study.

## Author contribution

SB, LY, and AG-M conceived the study and designed the experiments. SB, RS, GS, MS, and LY generated, curated, and performed data analysis. SB, GE, KK, and AG-M performed the physiological and molecular experiments. SB, LY, and AG-M wrote and edited the original manuscript. All authors discussed the results and contributed to the final version of the manuscript.

## Acknowledgements

We acknowledge the Greenhouse facility at CRAG for technical assistance. We thank Dr Víctor Manuel González (CRAG) for assistance in data analysis. Computational resources were provided by the e-INFRA CZ project (ID:90254) of the Czech Republic. We are grateful to Charlotte Poschenrieder for her mentorship in abiotic stress plant physiology. We acknowledge the *Catalan Initiative for the Earth BioGenome Project* for the inclusion of the species used in this study within its initiative.

## Data Availability

Sequencing data have been deposited in the European Nucleotide Archive under project accession PRJEB107246. Transcriptomic data are available at GEO under accession GSE309205, accessible to reviewers using the token ifgloeugfvcrdgh.

## Supplementary Information

**Supplementary Methods.** *Brassica fruticulosa* (**1**) genome sequencing and genome size estimation; (**2**) genome assembly, scaffolding, and quality assessment; (**3**) repeat identification and masking; and (**4**) evidence-guided gene prediction and functional validation.

**Figure S1.**
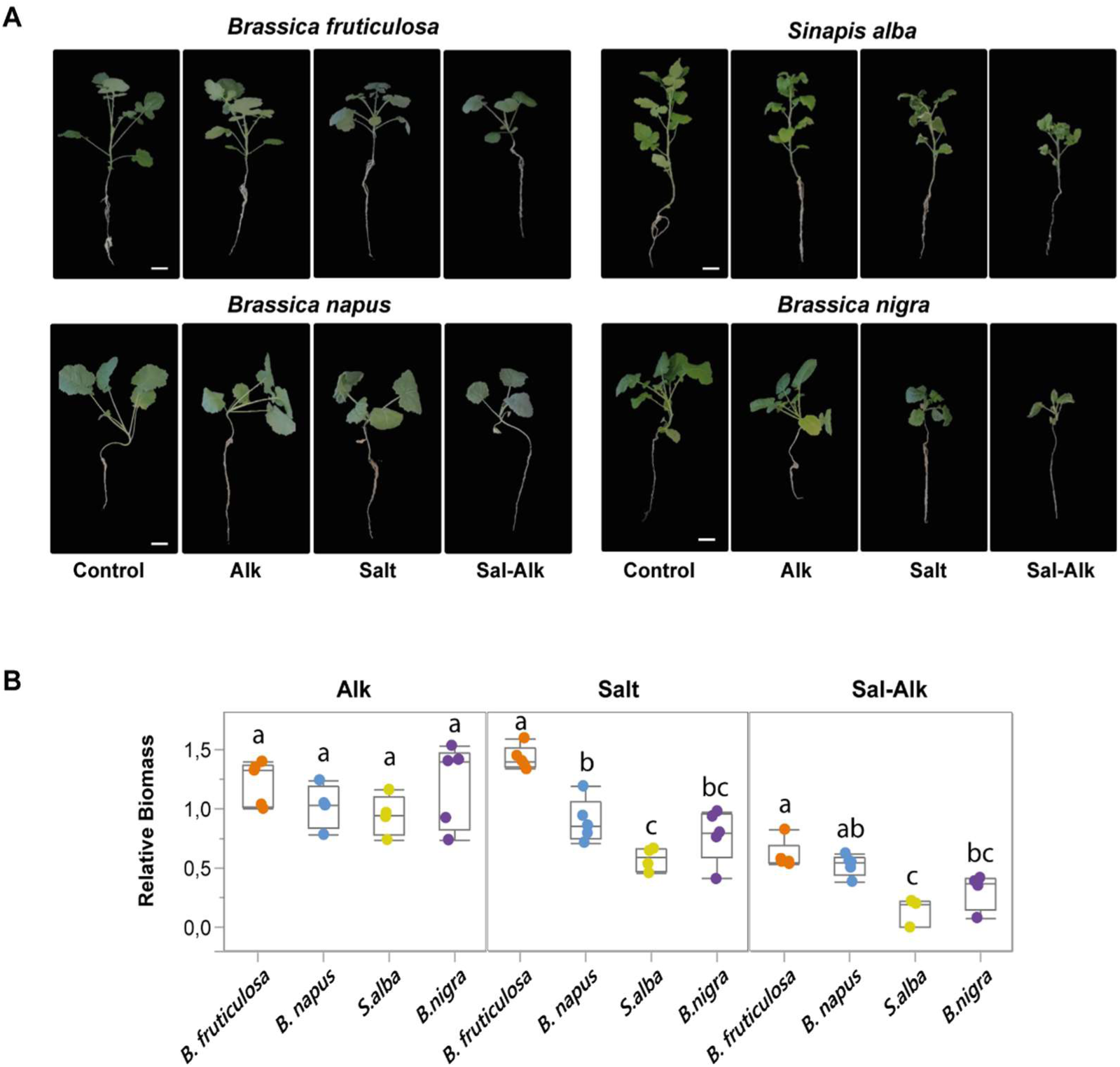
**A)** Representative pictures, and **B)** relative biomass (DWtreatment/control) of 25d old *B. fruticulosa*, *B. napus*, *S. alba,* and *B. nigra* plants cultivated under Control (½HS, pH 5.9), Alk (½HS + 15 mM NaHCO_3_, pH 8.3), Salt (½HS + 150 mM NaCl, pH 5.9), and Sal-Alk (½HS + 15 mM NaHCO_3_ + 135 mM NaCl, pH 8.3) conditions for 10 days. Letters indicate significant differences between species for each treatment (adjusted *P* ≤ 0.05, Tukey’s HSD test).

**Figure S2.**
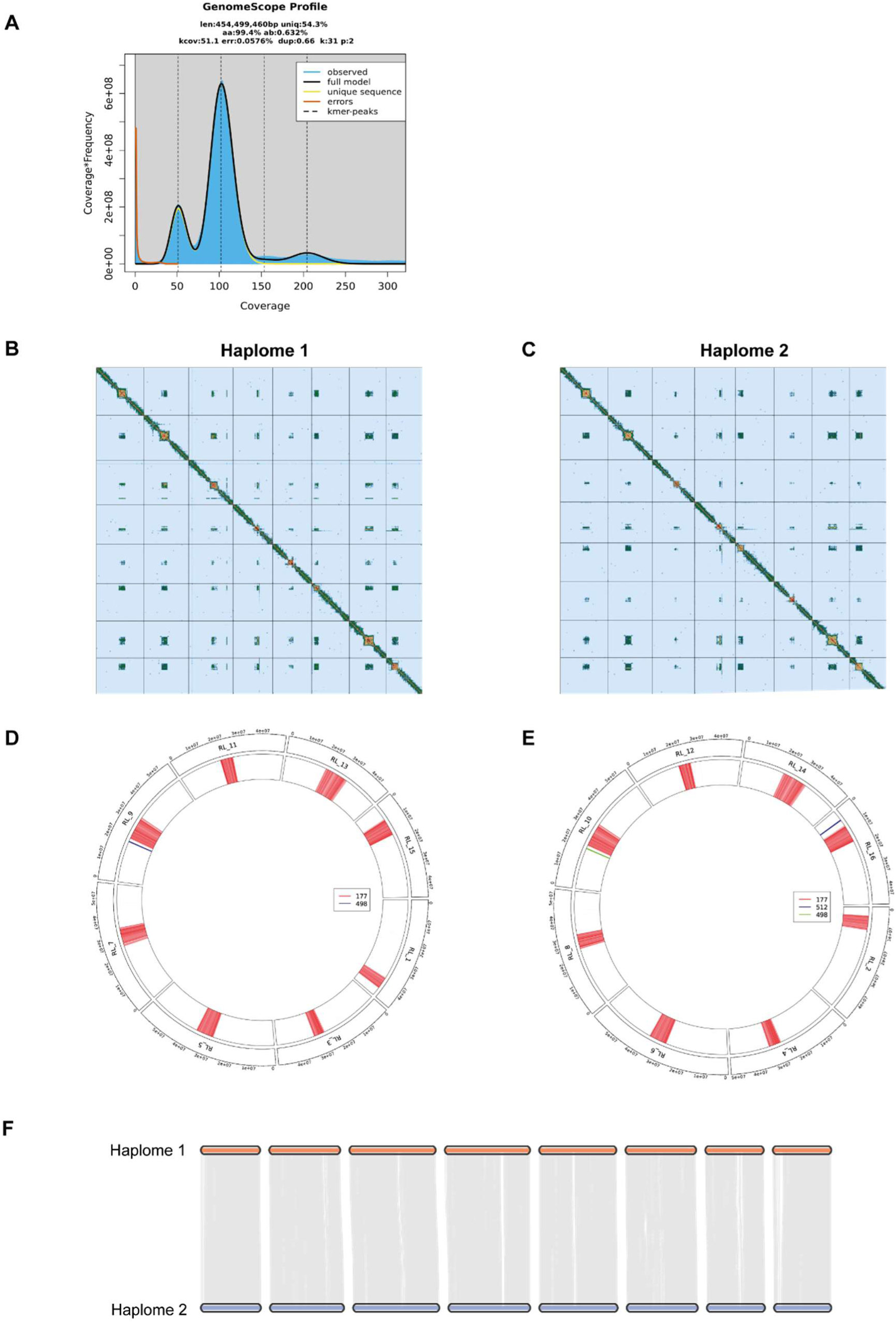
Concordant, chromosome-level dual haplome assemblies of *Brassica fruticulosa*. **A**) GenomeScrop2.0 kmer-based profile of B.fruticulosa genome. **B&C**) Hi-C chromosomal scaffolding contact frequency heatmaps. Blue to yellow to red indicates increasingly stronger Hi-C interactions. **D&E**) Centromere repeat landscape of both B. fruticulosa haplomes. These independently scaffolded genomes exhibit very high collinearity. **F**) Chromosome-level view of synteny between two *B. fruticulosa* haplomes.

**Dataset S1**. Transcriptome changes in *Brassica fruticulosa*.

**Dataset S2**. Transcriptome changes in *Brassica napus*.

**Dataset S3**. Transcriptome changes in *Sinapis alba*.

**Dataset S4**. Transcriptome changes in *Brassica nigra*.

**Dataset S5**. Arabidopsis ortholog conversion of transcripts of Brassicaceae species.

**Dataset S6**. Functional annotation of genes included in the heatmap for transcriptome changes.

**Dataset S7**. Gene Regulatory Networks.

**Dataset S8**. Orthologues for salt- and osmotic stress-responsive genes in distinct Brassicaceae species.

**Dataset S9**. Enrichment of Gene Regulatory Networks in transcription families.

**Dataset S10**. Potential cis-regulatory elements used in this study.

**Dataset S11**. List of additional Brassicaceae species used in this study.

**Dataset S12.** Hub transcription factors from each species’ GRNs.

## Supplementary Methods

### SM1. *Brassica fruticulosa* genome sequencing and genome size estimation

High molecular weight (HMW) DNA was extracted from a young seedling of *B. fruticulosa* accession ‘ESC’ using a modified CTAB protocol (Doyle & Doyle, 1987). DNA integrity was confirmed by gel electrophoresis and sheared to 15–20 kb. PacBio HiFi libraries were prepared and sequenced by Novogene (UK) Company Limited on one SMRT cell of the PacBio Sequel Revio system in CCS mode.

From the same individual, a Hi-C library was prepared from leaf tissue using the Arima Genomics® Hi-C kit and sequenced on an Illumina NovaSeq 6000 S4 platform (PE150). Plants were grown under natural glasshouse conditions in pots containing natural soil.

Genome size was estimated using k-mer (k = 31) distribution analysis with KMC (Kokot et al., 2017) and *GenomeScope v2.0* (Ranallo-Benavidez et al., 2020), with parameters: ploidy = 2, k-mer length = 31, and maximum k-mer coverage = 200,000. Independent genome size estimates were obtained by flow cytometry following Doležel et al. (2007).

### SM2. *Brassica fruticulosa* genome assembly, scaffolding, and quality assessment

Initial contig-level assemblies were generated using *hifiasm v0.24* (Cheng et al., 2021), integrating HiFi and Hi-C reads. Hi-C data were mapped to each haplotype assembly and processed following the Dovetail Omni-C pipeline. Scaffolding was performed with *YaHs v1.2.2* (Zhou et al., 2023).

Potential contamination was assessed using *BlobTools v4* (Laetsch & Blaxter, 2017), incorporating gene completeness metrics from *Compleasm v0.2.6* (Huang & Li, 2023), sequence coverage estimates generated with *Minimap* (Li, 2018), and taxonomic assignments based on *Diamond BLAST* (Buchfink et al., 2021). Contaminant contigs were removed prior to manual curation.

Assemblies were visualised using *PretextMap* and manually curated using *PretextView v0.2.5* following the rapid curation pipeline. Telomeric repeats were annotated using *tidk v0.2.1* (Brown et al., 2025), and coverage tracks were generated with *bedtools genomecov v2.3.1.1* (Quinlan & Hall, 2010). Centromeric regions were validated using *TRASH v1.2* (Wlodzimierz et al., 2023). Global assembly statistics were computed with *gfastats v1.3.1* (Cabanettes & Klopp, 2018), and assembly completeness was assessed using *Compleasm BUSCO* analyses (Shen et al., 2023).

### SM3. Repeat identification and masking

Repetitive elements were identified using a combination of de novo and homology-based approaches. A custom repeat library was constructed using *HelitronScanner v1.1* (Xiong et al., 2014), *LTR_FINDER v1.3* (Xu & Wang, 2007), *LTR_retriever v3.0.1* (Ou & Jiang, 2018), *MiteFinderII v1.0.006* (Hu et al., 2018), *RepeatModeler v2.0.6* (Flynn et al., 2020), and *SINE_Scan v1.01* (Mao & Wang, 2017). Predicted repeat elements were classified using *RepeatClassifier v2.0.6* (Flynn et al., 2020) and merged with Brassicaceae-associated repeats from the *Dfam v3.9 database* (Storer et al., 2021) and *RepBase v20181026* (Bao et al., 2015). The combined repeat library was clustered using *vsearch v2.30.0* (Rognes et al., 2016) to generate a non-redundant repeat dataset. The final repeat library was then used to mask the *B. fruticulosa* genome using RepeatMasker v4.1.8 (Flynn et al., 2020).

### SM4. Evidence-guided gene prediction and annotation

Protein-coding genes were predicted on the repeat-masked genome using an evidence-guided strategy integrating ab initio, homology-based, and transcriptome-based approaches. Ab initio gene prediction was performed using *BRAKER3 v3.0.8* (Gabriel et al., 2024), which automatically trained *Augustus v3.5.0* (Stanke et al., 2004) and *GeneMark-ETP v1.02* (Brůna et al., 2024) using RNA-seq alignments and reference protein sequences from the OrthoDB12 Viridiplantae dataset (Tegenfeldt et al., 2025).

RNA-seq data from 48 *B. fruticulosa* samples (European Nucleotide Archive project accession PRJEB74663) were quality-filtered using *fastp v0.20.148* (Chen et al., 2018) and aligned to the genome using *HISAT2 v2.2.1* (Kim et al., 2015). For homology-based prediction, *GeMoMa v1.9* (Keilwagen et al., 2019) was used with protein sequences from seven related Brassicaceae species: *Arabidopsis thaliana* (GCF_000001735.4), *Brassica nigra* (GCA_964214035.1), *Brassica oleracea* (GCF_000695525.1), *Brassica rapa* (GCF_000309985.2), *Hirschfeldia incana* (GCA_044115375.1), *Raphanus sativus* (GCF_000801105.2), and *Sinapis alba* (GWHBFWS00000000).

For transcriptome-based annotation, RNA-seq reads were assembled de novo using *Trinity v2.15.2* (Grabherr et al., 2011), followed by gene structure annotation with the *PASA pipeline v2.5.3* (Haas et al., 2003) to identify full-length transcripts. In parallel, reference-guided transcriptome assemblies were generated using *StringTie v2.2.1* (Kovaka et al., 2019), and coding sequences were predicted using *TD2 v1.0.5* (Mao et al., 2025).

Gene models generated by all three approaches were integrated using *EvidenceModeler v2.1.0* (Haas et al., 2008) and further refined with *PASA* (Campbell et al., 2006) to add untranslated regions and alternative splicing isoforms. Final gene models were filtered to remove entries with internal stop codons, missing introns, or duplicated features using *Webin-CLI* and *table2asn* (ENA and NCBI validation tools), ensuring compliance with ENA and NCBI annotation standards. Annotation completeness was evaluated using *BUSCO v5.8.2* (Manni et al., 2021) and *OMArk v0.3.0* (Nevers et al., 2025), employing the brassicales_odb12 dataset and the LUCA.h5 database.

